# A novel and highly specific Forssman antigen-binding protein from sheep polyomavirus

**DOI:** 10.1101/2023.04.10.536218

**Authors:** Nils H. Rustmeier, Lisete M. Silva, Antonio Di Maio, Joshua C. Müller, Alexander Herrmann, Ten Feizi, Yan Liu, Thilo Stehle

**Affiliations:** Interfaculty Institute of Biochemistry, University of Tübingen, Tübingen, Germany; Glycosciences Laboratory, Faculty of Medicine, Imperial College London, London, UK

**Author notes:** Corresponding author: Thilo Stehle.

## Abstract

Polyomaviruses are small, non-enveloped double-stranded DNA viruses of humans and other mammals, birds, and fish. Infections are usually asymptomatic and result in latency, however, some polyomaviruses can induce severe diseases, including cancer, in immunocompromised individuals. Established cellular receptors for polyomavirus infection are sialylated glycolipids (such as gangliosides), membrane proteins, and glycosaminoglycans. Polyomaviruses are usually highly host specific but the exact principles that govern host tropism remain unknown in many cases. Here, glycan array screening shows that the major capsid protein VP1 of sheep polyomavirus (ShPyV) binds to the Forssman glycolipid, an antigen of many vertebrates and a potential tumor marker in humans. Following closer investigation, we can report for the first time that a neutral, non-sialylated glycolipid acts as a polyomavirus receptor. Concurrently, we present the first report of a viral protein that specifically engages the Forssman antigen. We demonstrate that ShPyV VP1 binds to Forssman-positive erythrocytes but not those of human A, B and O blood groups, which is a clear distinction from features thus far described for Forssman lectins. X-ray crystallography and structural analysis of the VP1-Forssman glycan complex define the terminal Forssman disaccharide as the determinant of this protein-receptor interaction. These results strongly suggest that the sheep polyomavirus can use Forssman antigen for infectious cell entry. Furthermore, the ability of ShPyV VP1 to distinguish Forssman-positive from -negative cells may prove useful for monitoring the Forssman-‘status’ of normal, preneoplastic and neoplastic cells and tissues and establishing the antigen level as a biomarker.

**Author summary:** Elucidation of host cell receptor specificities of viral infection is crucial to understand the pathobiology of associated diseases and develop treatments. However, for many polyomaviruses the receptor engagement as the initial event in infection is poorly understood. In only a few cases polyomavirus tropism has been pinned down to a single type of glycan receptor. While many polyomaviruses utilize sialyl glycans to attach to host cells, the role of non-sialylated glycans as receptors is so far underestimated. Here, we show for the first time that a glycan of neutral charge, in this case the carbohydrate portion of the Forssman antigen, acts as a ligand for a polyomavirus capsid protein and may thus contribute to host tropism and infective cell entry. These results represent a significant addition to knowledge on polyomavirus-glycan interactions and complement general principles of carbohydrate engagement by viruses. Furthermore, as a specific binding protein of Forssman antigen, VP1 may help to determine levels of this antigen in healthy and malignant tissues in humans.

## Introduction

The Forssman antigen (FA), a neutral globo-series glycosphingolipid (GSL, see Table 1), is produced by the glycosyltransferase activity of Forssman synthetase (FS)^1, 2^. Encoded by the *GBGT1* gene, FS catalyzes the addition of αGalNAc to the non-reducing end of globoside (see Table 1). This reaction results in the key immunogenic motif of the Forssman antigen, the terminal GalNAcα1-3GalNAcβ Forssman disaccharide^3^. With few exceptions in humans, *GBGT1* is transcribed but encodes for an inactive FS, resulting in a predominantly Forssman-negative human population^4, 5^. So far, only a small number of healthy individuals worldwide were found to carry an activating mutation within *GBGT1.* In these individuals, the resulting allele gives rise to a Forssman antigen-positive (FA+) erythrocyte phenotype, which is now recognized as the human FORS blood group^6–8^. Other reports of healthy human FA+ tissues are scarce, and human sera often contain natural anti-FA antibodies^9–11^. More frequently, Forssman antigenicity was described in cancerogenesis and a few other diseases of humans^12–21^. Yet, both the genetics and the precise role of FA expression in disease remain poorly understood.

**Table 1.**
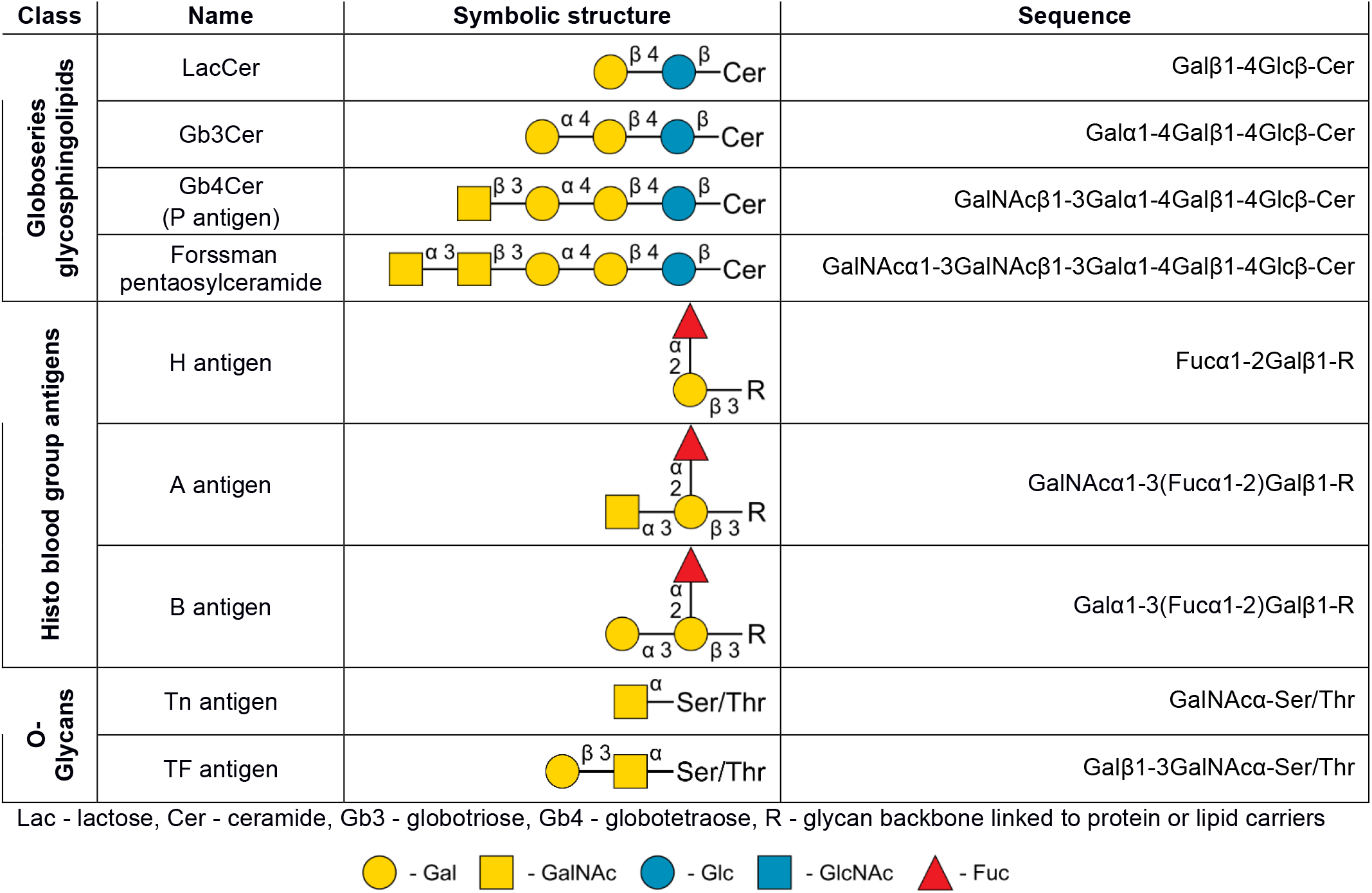
Carbohydrate antigens cited in this report.

| Class | Name | Symbolic structure | Sequence |
| --- | --- | --- | --- |
| Globoseries glycosphingolipids | LacCer | | Galβ1-4Glcβ-Cer |
| | Gb3Cer | | Galα1-4Galβ1-4Glcβ-Cer |
| | Gb4Cer (P antigen) | | GalNAcβ1-3Galα1-4Galβ1-4Glcβ-Cer |
| | Forssman pentaosylceramide | | GalNAcα1-3GalNAcβ1-3Galα1-4Galβ1-4Glcβ-Cer |
| Histo blood group antigens | H antigen | | Fucα1-2Galβ1-R |
| | A antigen | | GalNAcα1-3(Fucα1-2)Galβ1-R |
| | B antigen | | Galα1-3(Fucα1-2)Galβ1-R |
| O-Glycans | Tn antigen | | GalNAcα-Ser/Thr |
| | TF antigen | | Galβ1-3GalNAcα-Ser/Thr |

Lac - lactose, Cer - ceramide, Gb3 - globotriose, Gb4 - globotetraose, R - glycan backbone linked to protein or lipid carriers
- Gal - GalNAc - Glc - GlcNAc - Fuc

The Forssman glycolipid also serves as a receptor for different bacterial infections in animals^22–28^, however, it is so far unknown if viruses also utilize FA to target their hosts. In this study, by means of glycan microarray screening, nuclear magnetic resonance (NMR), surface plasmon resonance (SPR), and cell binding assays, we show that FA, which is the predominant glycolipid in ovine erythrocytes^29^, serves as a cellular attachment receptor for the VP1 of a polyomavirus originally identified in sheep^30^. This designated sheep polyomavirus (ShPyV) is a member of a non-enveloped, double-stranded DNA virus family with icosahedral capsids, scaffolded by the major capsid protein VP1. To initiate an infection many polyomaviruses rely on the interaction between their VP1 and sialic acid-equipped glycolipids in the target cell membranes^31–,36^. Although other infection receptors such as glycoproteins and glycosaminoglycans have also been reported for some polyomaviruses^37–42^, detailed structural information is so far only available for the interactions between VP1s and sialylated oligosaccharides. Thus, our studies present the first example of a polyomavirus VP1 protein interacting with a non-sialyl neutral glycan, complementing the principles of how polyomaviruses recognize their targets.

## Results

### Sheep polyomavirus VP1 binds the Forssman epitope with a strong preference over other glycans

To identify potential attachment factors of ShPyV, we prepared recombinant VP1 pentamers as previously described^43^. The protein was analyzed using a neoglycolipid-based glycan screening array encompassing a broad spectrum of lipid-linked glycan probes^44^. A full list of 672 glycan probes is in Table S1, and these include major types of mammalian sequences found on glycoproteins (N-and O-linked), glycolipids, blood group and Lewis antigen related, oligosaccharide fragments of glycosaminoglycan heparin, as well as those derived from polysaccharides of bacteria, fungi, and plants.

Among these are 204 sialylated glycan probes variously bound by several polyomavirus VP1s described previously^45–49^. We detected a signal for the Forssman glycolipid that surpasses all other signals by at least one order of magnitude (Fig. 1). A dominance of the terminal GalNAcα1-3GalNAcβ Forssman disaccharide is apparent as the VP1 did not give binding signals with globoside glycolipid (P-antigen, position 25, Table S1), the precursor of the Forssman glycolipid which lacks the terminal α-linked GalNAc residue. Furthermore, there were no signals with other GalNAc-terminating glycan probes, including the blood group A related glycan probes, and the two Tn antigen probes GalNAcα-Ser and -Thr (positions 221 and 222, Table S1). The disaccharide probe GalNAcα1-3GalNAc (position 223) with a ring-opened GalNAc core (reduced by reductive amination) was also not bound. These suggest the importance of adjoining monosaccharides beyond the αGalNAc residue for ShPyV VP1 binding. Among the sialylated glycans analyzed only one short α2,6-linked sialyl probe with a 9-O-acetylated N-acetylneuraminic acid residue (Neu5,9Ac_2_) yielded a weak signal (position 607). No binding signals were detected in the negative control experiment with detection antibodies alone in the absence of ShPyV VP1 (Table S1).

**Figure 1.**
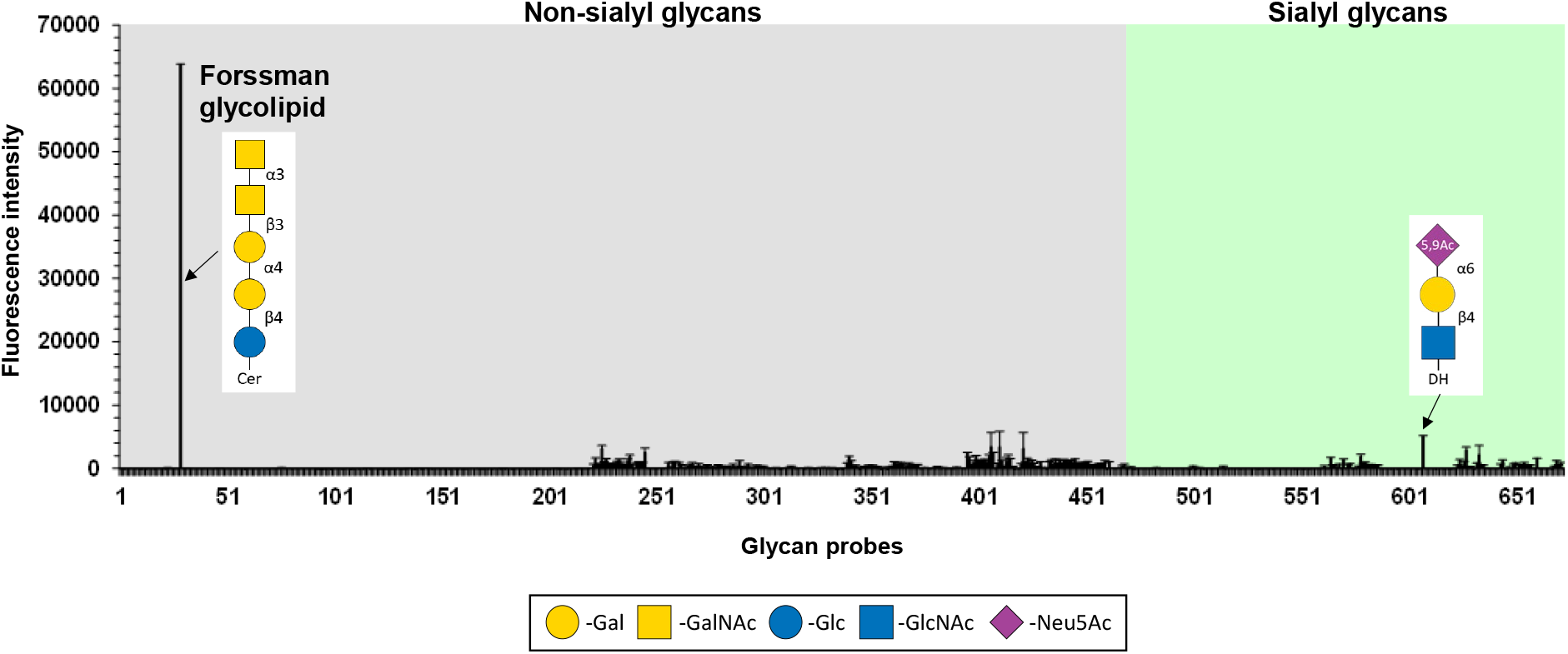
Glycan microarray screening analysis of ShPyV VP1 reveals selective engagement of Forssman glycolipid. The results are the means of fluorescence intensities of duplicate spots, printed at 5 fmol per spot. The error bars represent half of the difference between the two values. In the glycan array the 672 lipid-linked probes are grouped into non-sialylated and sialylated glycans as annotated by the colored panels. The list of glycan probes, their sequences, and binding scores are in Table S1.

To assess if the glycan binding revealed by the glycan microarray also occurs in solution, we evaluated the interactions between ShPyV VP1 and the Forssman pentaose (F_P_) in the presence of α2,6-sialyllactose (6’SL) by saturation transfer difference NMR (STD NMR) spectroscopy, which reveals protein-to-ligand magnetization transfer in the case of short-term binding (10^-8^ M < K_D_ < 10^-3^ M)^50, 51^. In the first instance, the ^1^H reference resonances of isolated F_P_ and 6’SL were collected (Fig. 2a and 2b), which allowed us to unambiguously assign the oligosaccharide signals^43, 52^ (except for the overlaps within 3.5-4.0 ppm) in the ^1^H spectrum of a sample comprising VP1, F_P_, and 6’SL (Fig. 2c). Subsequently, we used the same sample to perform the STD NMR experiment. Evaluation of the resulting spectrum reveals magnetization transfer to both F_P_ and 6’SL, which infers that VP1 interacts with both glycans in the solution (Fig. 2b). However, while resonances derived from all monosaccharides in F_P_ (two GalNAc, two Gal, and a Glc) received magnetization transfer, only the sialic acid methyl group of 6’SL re-emerged in the STD spectrum. Yet, these results are in good agreement with the findings from the glycan array analysis of ShPyV VP1, which indicated a much higher affinity for F_P_ than sialylated glycans (Fig. 1). Strikingly, the methyl group of the αGalNAc in F_P_ gives a particularly strong peak in the STD spectrum (Fig. 2b), which implies a prominent (and potentially specificity-conferring) role in the VP1-F_P_ interaction.

**Figure 2.**
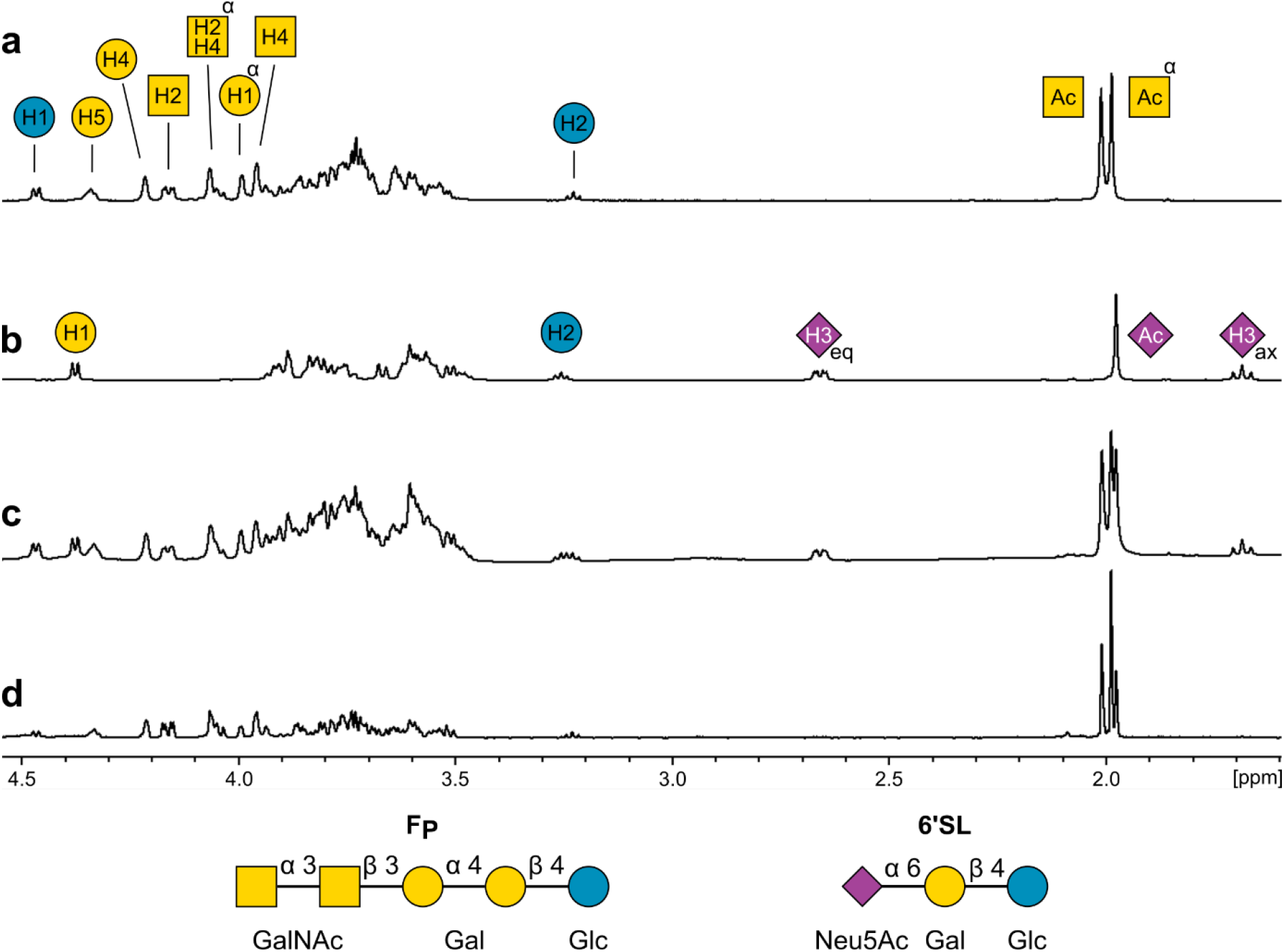
Assessment of the sheep polyomavirus VP1 – Forssman pentaose interaction in solution by STD NMR. ^1^H spectra of **a** Forssman pentaose (F_P_) and **b** 6’sialyllactose (6’SL) in their isolated forms. Proton resonances are labeled according to the carbon positions in the monosaccharides. **c** _1_H spectrum of a sample comprising 20 µM ShPyV VP1, 1 mM F_P_, and 1 mM 6’SL collected under protein suppression. **d** The STD NMR spectrum that was recorded from the same sample.

### ShPyV VP1 has a nanomolar affinity towards Forssman pentaose-decorated surfaces

After assessing the complex formation between ShPyV VP1 and F_P_ in the context of the solid-supported glycan array and solution NMR, we sought to evaluate the underlying binding kinetics by surface plasmon resonance (SPR) and compare the performance of VP1 with the albumen gland agglutinin of *Helix pomatia* (HPA), an αGalNAc-specific lectin with high affinity towards the Forssman antigen^53, 54^. Therefore, we prepared an SPR chip decorated with F_P_ (see Fig. S3) and measured binding responses using different dilutions of VP1 and HPA. The resulting response curves show that within the given association time the pentavalent VP1 and hexavalent HPA bind to the immobilized F_P_ without reaching a complex formation equilibrium, which suggests that the proteins do not readily dissociate from the surface (Fig. 3a). Indeed, fitting of the kinetic parameters results in extremely low dissociation rates (k_off_) for both proteins, which is a common phenomenon for multivalent binding systems. However, the curves of VP1 result in a higher rate for complex formation (k_on_), determining an overall approximately three-fold lower dissociation constant (K_D_) compared to HPA (Fig. 3a).

**Figure 3.**
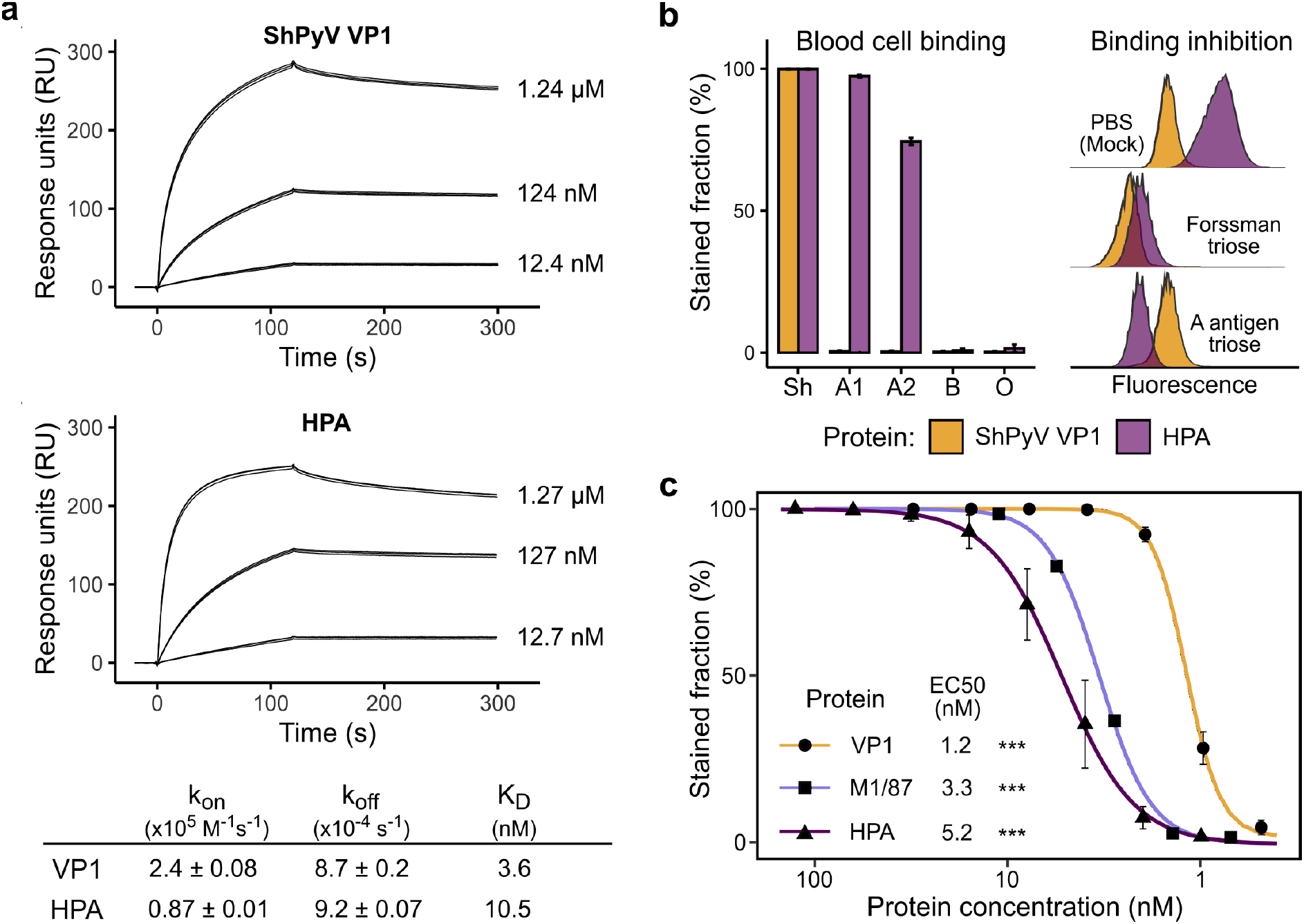
Flow cytometry and surface plasmon resonance analyses of Forssman antigen engagement. **a** Determination of the binding kinetics towards a Forssman-decorated surface using ShPyV VP1 and HPA. Response curves are shown as triplicates for each protein concentration. Kinetic parameters and the dissociation constant are provided in the table below. **b** Absolute cell attachment of ShPyV VP1 and *Helix pomatia* agglutinin (HPA) to Forssman antigen-positive red blood cells (RBCs) from sheep (Sh) and human erythrocytes of the blood groups A1, A2, B, and O (left). Effects of Forssman and blood group A antigen trioses on ShPyV VP1 and HPA attachment to sheep RBCs. Results are shown as concatenated and normalized histograms (right). **c** EC50 determination using ShPyV VP1, HPA, and anti-Forssman IgM M1/87 to sheep blood cells. Significance is indicated with *** for p < 0.001.

### ShPyV VP1 specifically binds to Forssman-positive erythrocytes

To confirm the functional relevance of the VP1-F_P_ complex *in vitro*, we performed cell binding assays using different erythrocytes and compare the cell binding of VP1 with commercial reagents for Forssman antigen detection, i.e., HPA and the monoclonal anti-FA IgM antibody M1/87^55^. We incubated VP1 with Forssman-positive red blood cells (RBCs) from sheep and human RBCs of the different ABO phenotypes and found that ShPyV VP1 bound sheep RBCs with saturation but did not attach to any of the human erythrocytes (Fig. 3b left). In contrast, the αGalNAc-specific lectin HPA bound to the FA on sheep RBCs and to the antigens on human blood group A1 and A2 RBCs^56^ (Fig. 3b left). To confirm that the carbohydrate portions of the antigens account for binding, we incubated the two proteins with excessive amounts of the terminal trisaccharides (trioses) of FA (GalNAcα1-3GalNAcβ1-3Gal) and A substance (GalNAcα1-3[Fucα1-2]Gal) and assessed the remaining sheep RBCs attachment. As predicted, FA triose blocked VP1 and HPA, while A antigen triose could only inhibit HPA attachment (Fig. 3b right). Lastly, we determined the effective protein concentrations (EC50, see Methods) for sheep RBCs attachment using different dilutions of ShPyV VP1, HPA, and M1/87. Evaluation of the response curves yielded the lowest EC50 for ShPyV VP1 with a value of 1.2 nM, followed by M1/87 and HPA with 3.3 nM and 5.2 nM, respectively (Fig. 3c). Collectively, these results show the remarkable specificity and affinity of ShPyV VP1 towards the Forssman antigen.

### Crystal structure of Forssman pentaose bound to VP1 reveals an intricate network of interactions

To characterize the binding of Forssman antigen by ShPyV VP1 with atomic detail, we solved the crystal structure of the VP1 in complex with 10 mM of Forssman pentaose (F_P_) at 1.92 Å resolution (Table S4). Initial electron density calculation using unliganded ShPyV VP1 model (PDB 6Y61) shows that five molecules of the oligosaccharide (GalNAcα1-3GalNAcβ1-3Galα1-4Galβ1-4Glc) associate with the outer surface of the pentameric VP1, at a position that does not overlap with the previously described sialic acid binding site^43^ (Fig. S5b), in a five-fold symmetric array. All five Forssman binding sites are identical and located between two neighboring VP1 monomers (Fig. 4a). In the binding sites without artificial crystal contacts, electron density provides an unambiguous outline of the last four monosaccharides of F_P_ (Fig. 4b). The intermolecular contacts show that the terminal trisaccharide of F_P_ contributes to binding, whereas the remaining βGal and Glc divert away from VP1 and do not interact with the protein surface. Closer inspection of the binding site reveals that the methyl group of the terminal αGalNAc invades a hydrophobic surface depression constituted by phenylalanine residues of two neighboring VP1 monomers, establishing van der Waals interactions (Fig. 4c). Additionally, the *N*-acetyl moiety of αGalNAc interacts with the Q266 and F66cw residues via hydrogen bonds (Fig. 4c). In βGalNAc, the carbonyl oxygen forms two hydrogen bonds with the side chains of Q266 and N268 (Fig. 4c). Apart from these contacts, numerous interactions involve the hydroxyl groups of F_P_ that are directed towards the protein surface. An overview of all contacts and their distances is provided in Table 2. We also solved the crystal structure of VP1 in complex with the FA glycan precursor, globo-*N*-tetraose, which reveals negligible binding only (Fig. S5). We therefore conclude that the terminal GalNAcα1-3GalNAc disaccharide is essential for the high affinity of the ShPyV VP1-F_P_ complex.

**Figure 4.**
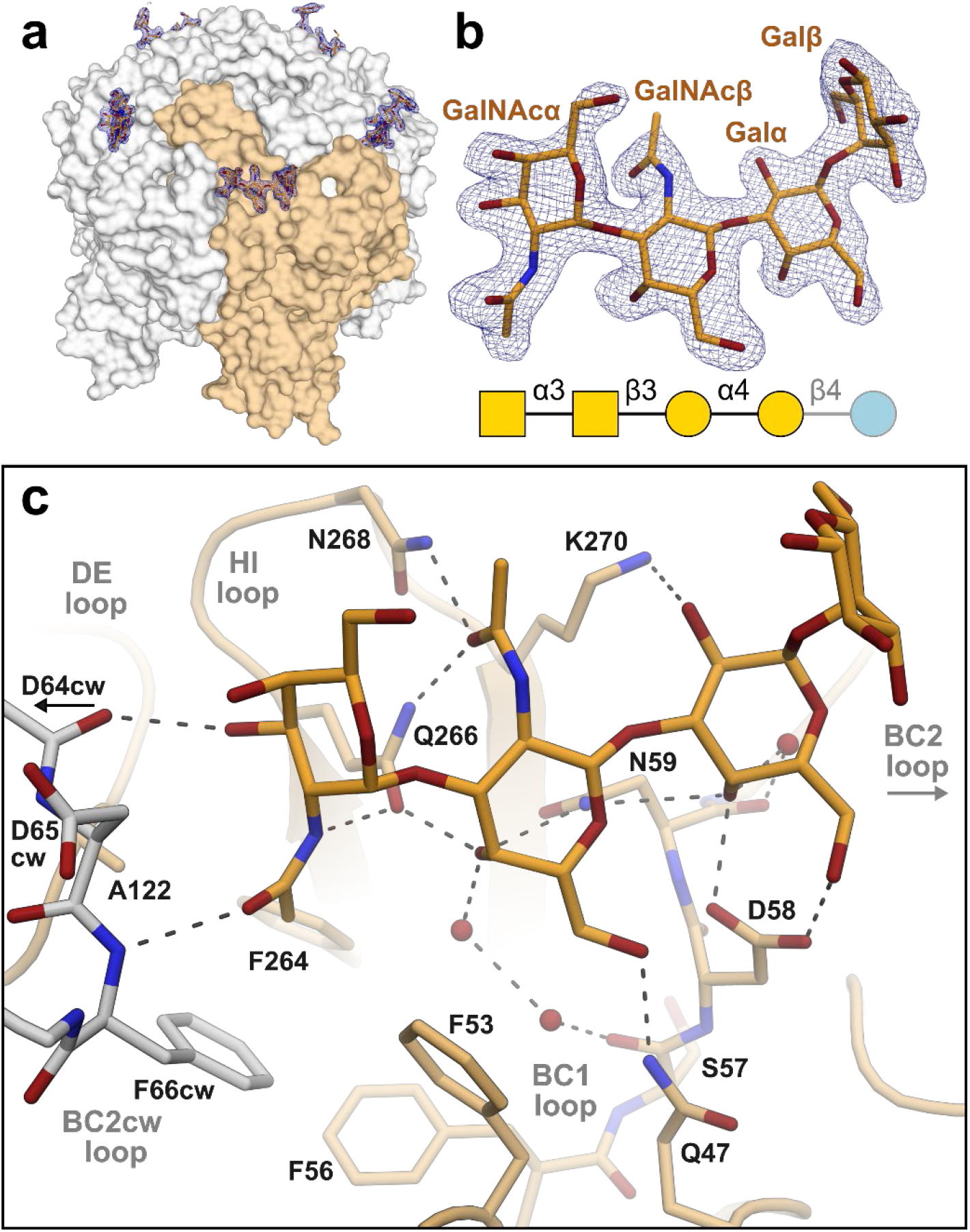
Forssman pentaose binds to the external surface of ShPyV VP1 with numerous contacts. **a** The crystal structure of a single VP1 pentamer is shown with five molecules of Forssman pentaose (F_P_) associated to its outer surface. The bias-reduced Fo-Fc omit electron density map of F_P_ is displayed as blue meshes around the oligosaccharides, with a contour level of 2.5 σ and a radius of 1.6 Å. The individual F_P_ binding sites locate between adjoining protein chains, indicated by the orange coloring of one VP1 monomer. **b** Magnified view of a bound F_P_ molecule with its symbol structure. The glucose of F_P_, which is not visible in the crystal structure, is grayed out in the latter. **c** Molecular interactions between VP1 and F_P_ are displayed with the hydrogen bonds drawn as dashed lines. VP1 coloring corresponds to panel **a**. Water molecules are shown as spheres. Throughout the figure, oxygen and nitrogen atoms are colored red and blue, respectively.

**Table 2.**
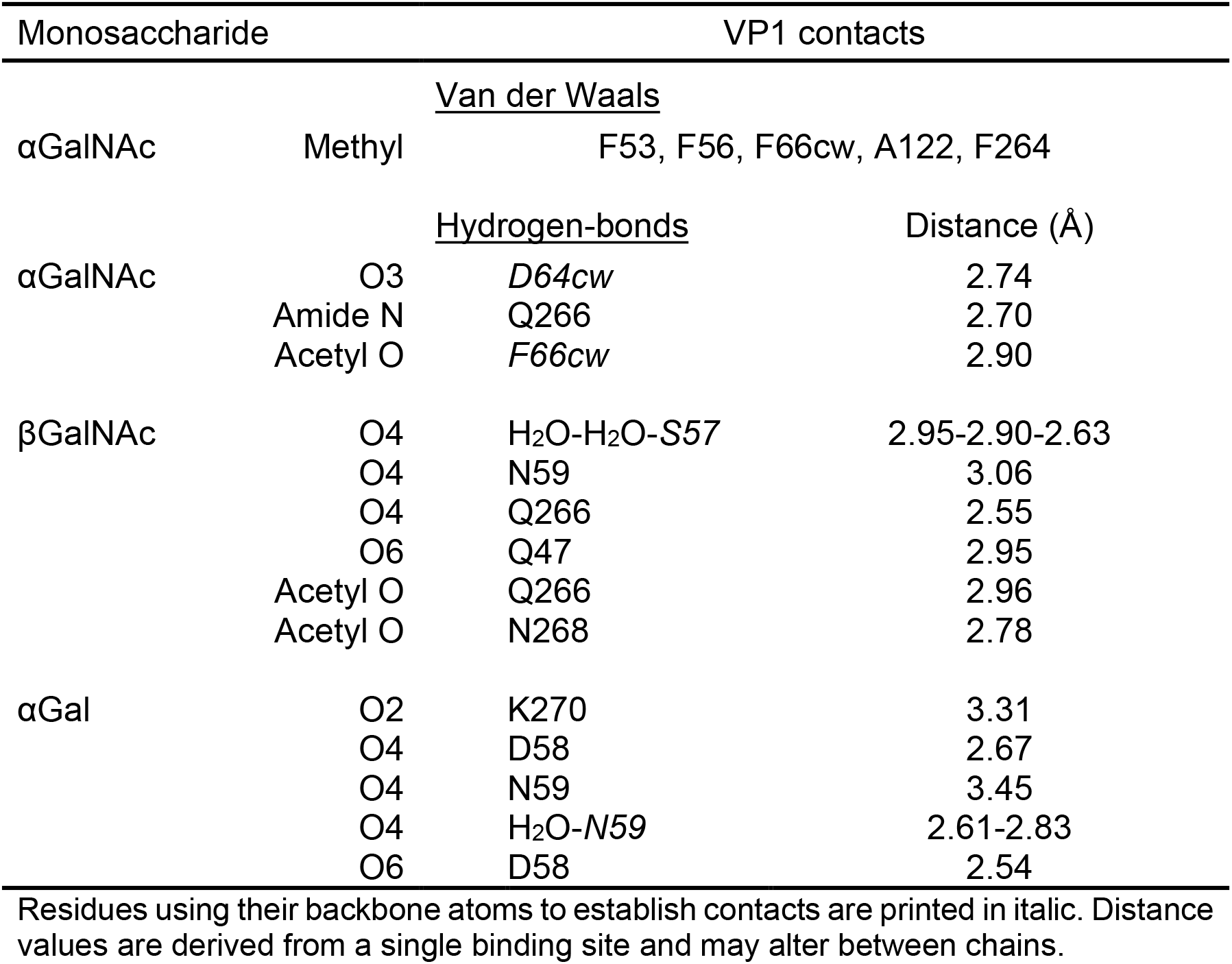
Intermolecular contacts between ShPyV VP1 and Forssman pentaose.

## Discussion

Sheep polyomavirus (ShPyV) was originally identified as a genomic contaminant in processed lamb meat, and it remains unknown if sheep are the natural hosts of this virus^30^. Our functional data show that ShPyV engages Forssman antigen (FA)-decorated cells with high affinity and specificity, making it plausible that FA initiates sheep polyomavirus infection. However, in sheep the FA was thus far only detected on erythrocytes but not on leukocytes, bone marrow or other tissues^29, 57, 58^. However, as the studies about the distribution of Forssman antigen in sheep tissues are old, the more sensitive techniques available today could help to elucidate the potential targets cells for viral replication as well as the precise role of Forssman antigen during ShPyV infection.

The antigenic determinant of Forssman antigen is the GalNAcα1-3GalNAcβ disaccharide. In sheep erythrocytes it is predominantly linked to globotriasolyceramide, giving rise to the linear Forssman pentaosylceramide (see Table 1), but it also appears on the (unbranched) glycans of neolacto-series or Galili glycolipids^29^. In other animals, GalNAcα1-3GalNAcβ occurs on branched structures such as extended H-like antigens (dog gastric mucosa and mucus)^59, 60^, GM1 ganglioside (English sole fish)^61^, isoglobo-ganglio-neolacto glycans (swine)^62^, as well as in the form of mucin core 5 (GalNAcα1-3[NeuAca2-6]GalNAc) in bovine submaxillary mucin^63^. We believe that ShPyV VP1 can bind a range of different Forssman glycans, providing that the structural deviations do not result in clashes with the protein. By the assessment of our ShPyV VP1-Forssman pentaose (F_P_) complex structure, the designated minimal glycan binding motif is the GalNAcα1-3GalNAc disaccharide with few additional contacts to the subsequent αGal. From structural perspective, this epitope recognition clearly disallows branching of the glycan at certain positions, e.g., O3 of αGalNAc, N2 of βGalNAc or O4 of αGal. For example, in the blood group A glycan fucose replaces the *N*-acetyl group of the βGalNAc, which would result in overlaps with the protein and accounts for the lack of binding (Fig. S5b). This feature distinguishes VP1 from many other Forssman glycan-binding proteins such as Galectin-9, *Helix pomatia* agglutinin, *Sinularia lochmodes* lectin-2, and *Dolichos biflorus* agglutinin, where recognition is based on the subterminal or terminal GalNAc (see Fig. S6), resulting in a much broader glycan binding capacity^64–67^.

Previously we have shown that ShPyV VP1 interacts with sialylated trisaccharides in crystal structures^43^. Also, our new results from STD NMR and crystallography indicate that sialyl and Forssman glycan binding occur independently from each at separate sites of the protein surface (Fig. S5b). Yet, in our flow cytometry assays we did not detect any binding to human erythrocytes although they are known to be decorated with sialic acids. However, the preferred form of sialic acid for ShPyV VP1, as evinced by our glycan array data, is 9-*O*-acetylated *N*-acetylneuraminic acid (Neu5,9Ac_2_), which in human erythrocytes is present in too low amounts to result in detectable binding^68^. These findings imply that human cells are incompatible targets for the sheep virus.

Humans are considered a largely Forssman-negative species and the presence of Forssman antigenicity is mostly associated with human cancer^69^. During the last decade, the elucidation of the genetic basis behind FA reactivation and the establishment of the human FORS blood group underline the continual relevance of Forssman antigen^5, 6^. We propose that ShPyV VP1 complements and expands the available resources for the comprehensive detection of Forssman antigen in cellular and histological samples of healthy or malignant provenance. We also anticipate refining mutant VP1 protein to precisely target the Forssman antigen for biomedical use.

## Summary

We show that the Forssman antigen serves as a cellular attachment receptor for the major capsid protein VP1 of ShPyV. After initial detection by glycan array screening, cell binding assays imply functional relevance. Structural analysis of VP1 in complex with the carbohydrate moiety of FA, Forssman pentaose (FP), characterizes the determinants of the underlying interactions with high accuracy. Our findings have significance for the fields of structural virology, glycobiology, and pathobiology as follows:

i. We report the first structural and functional data of a viral entity to specifically interact with the Forssman antigen.
ii. The ShPyV VP1 – F_P_ complex provides the first detailed information on how a polyomavirus binds to a non-sialylated ligand.
iii. The high specificity and affinity of ShPyV VP1 toward Forssman antigen recommends this protein as a potential biomedical reagent.

## Materials and methods

### Protein expression and purification

The VP1 of sheep polyomavirus was recombinantly produced as previously.^43^ Briefly, transformed *E. coli* BL21 (DE3) were grown in LB (Miller) medium supplemented with 100 µg/ml ampicillin at 37 °C. Expression was induced at an OD_600_ of 0.7 with 400 µM IPTG and carried out at 20 °C for 16 h. ShPyV VP1 was extracted from bacterial pellets by sonification and subsequent centrifugation (17,000 rpm at 4 °C). The His_6_-tagged VP1 was purified by immobilized-metal affinity chromatography (IMAC) and gel filtration (20 mM HEPES pH 7.5, 150 mM NaCl). For crystallization of ShPyV VP1 the His_6_-tag was cleaved off by incubation with 10 U/mg thrombin overnight at room temperature. Tags and thrombin were separated from ShPyV VP1 by IMAC and gel filtration. If not immediately used, the protein was stored at -20 °C.

### Glycan array

The His_6_-tagged ShPyV VP1 was analyzed in the neoglycolipid (NGL)-based microarray system^70^. A broad-spectrum screening glycan microarray set encompassing 672 sequence-defined lipid-linked glycan probes was used (the glycan probes included, and their sequences are given in Supplemental Table S1). Details of the preparation of the glycan probes and the generation of the microarrays are in Supplementary Glycan Microarray Document (Supplemental Table S2) in accordance with the MIRAGE (Minimum Information Required for A Glycomics Experiment) guidelines for reporting of glycan microarray-based data^71^. The microarray analyses were performed essentially as described^47, 48^. In brief, after blocking the slides for 1h with HBS buffer (10 mM HEPES, pH 7.4, 150 mM NaCl) containing 0.33% (w/v) blocker Casein (Pierce), 0.3% (w/v) Bovine Serum Albumin (Sigma-Aldrich) and 5 mM CaCl_2_, the microarray was overlaid with the VP1 protein for 90 minutes as a protein-antibody complex that was prepared by preincubating VP1 with mouse monoclonal anti-polyhistidine and biotinylated anti-mouse IgG antibodies (both from Sigma) at a ratio of 4:2:1 (by weight) and diluted in the blocking solution to provide a final VP1 concentration of 150 µg/mL. Binding was detected with Alexa Fluor-647-labelled streptavidin (Molecular Probes) at 1 µg/mL for 30 minutes. Apart from the protein-antibody precomplexation step, which was performed on ice, all other steps were carried out at ambient temperature. Microarray imaging and data analysis are described in the Supplementary MIRAGE document (Supplemental Table S2).

### NMR

All nuclear magnetic resonance (NMR) data were collected in a Bruker AVIII-600 MHz spectrometer equipped with a TXI-z probe head at 298 K using 3 mm NMR tubes. Samples were provided in 99 % D_2_O-PBS containing 20 mM KH_2_PO_4_/K_2_HPO_4_ (pH 7.4) and 150 mM NaCl. ^1^H reference spectra of isolate oligosaccharides samples were recorded using pure solutions of 4 mM of Forssman pentaose and 2 mM of 6’SL. Subsequently, the ^1^H oligosaccharide resonances of a sample containing 1 mM Forssman pentaose and 1 mM 6’SL in the presence of 20 µM ShPyV VP1 were collected under suppression of protein signals. From the same sample, the saturation difference transfer (STD) NMR experiment was performed using previously published pulse protocols^43^. NMR spectra were processed using TOPSPIN 4 (Bruker). The carbohydrates used here were purchased from Biosynth, UK.

### VP1 mutagenesis

Site-directed mutagenesis of VP1 was performed in samples of 20 µl containing 1x ReproFast amplification buffer (Genaxxon), 20 ng template vector DNA, 50 ng primers, 2 mM dNTP, and 1 unit of ReproFast polymerase (Genaxxon). Initial denaturation was performed at 95 °C for 2 minutes, followed by 18 cycles of 95 °C for 1 minute (denaturation), 55-70 °C for 1 minute (annealing), and 72 °C for 6 minutes (elongation). Subsequently, DNA synthesis was completed at 72 °C for 10 minutes. Parental template DNA was digested with 2 units of DpnI for two hours at 37 °C. The PCR product was directly transformed into *E. coli* DH5α.

DNA primers for the site-directed mutagenesis of ShPyV VP1 were ordered from ThermoFisher Scientific and are displayed from 5’-to 3’ ends:

S95C_fwd: GCTGAACCAAGACATGACCTGCGATACCATCCTGATGTGGGAGG

S95C_rev: CCTCCCACATCAGGATGGTATCGCAGGTCATGTCTTGGTTCAGC

### Fluorescence labeling

VP1 was expressed and purified as described above. Surficial cysteine residues were reduced in degassed gel filtration buffer (20 mM HEPES pH 7.4, 150 mM NaCl) supplemented with 80 mM DTT at 4 °C overnight. DTT was removed from the protein solution using a PD-10 desalting column (Cytiva) with DTT-free gel filtration buffer. A 20-fold molar excess of cysteine reactive AFDye 488 Maleimide (Fluoroprobes) was added to the protein solutions (0.5 -0.7 mg/ml) and the reaction was incubated in the dark at 4 °C for 16-20 h. Excess dye was quenched and removed by desalting in gel filtration buffer supplemented with 10 mM DTT. VP1-488 conjugate was eluted in the same buffer. The degree of labeling was photometrically determined according to the manufacturer’s protocol (Fluoroprobes).

### Flow cytometry

Defibrinated blood from sheep was purchased from TCS Biosciences Ltd and BioTrading Benelux B.V., and human reference erythrocytes from Immucor. Prior to analysis, all blood was diluted to 0.25 % in 1x PBS + 2 % fetal calf serum (FC buffer). For staining, labeled protein was added to samples of ∼1×10^6^ cells in a total volume of 100 µL. After incubation in the dark for 15 minutes, cells were precipitated by centrifugation (1.5 min at 1900 rpm) and resuspended in fresh FC buffer. Unstained reference samples were prepared accordingly without the addition of labeled protein. For cell attachment inhibition assays, proteins were incubated with oligosaccharides (Forssman antigen trisaccharide: Elicityl, France; Blood group A trisaccharide: Biosynth, UK) for 5 minutes prior to their addition to the cellular samples (∼1×10^6^ cells in 100 µL). To determine the concentration for half maximal cell attachment (effective concentration, EC50), samples comprising serially diluted concentrations of labeled protein, starting from 10 µg/mL for M1/87-FITC (Santa Cruz Biotechnology) and HPA Alexa Fluor^TM^ 488 conjugate (Invitrogen), and 5 µg/mL for VP1-488, were prepared as described above. Here, the wash step was omitted to guarantee consistent protein concentrations and to prevent agglutination. All samples were transferred to the flow cytometer (LSR II, BD Biosciences) and fluorescent fractions were captured at 520 nM. 100,000 events were recorded for all samples except those with agglutination, in which case 10,000-20,000 events were recorded. All data were analyzed using FACSdiva software (BD Biosciences). EC50 values were calculated using R^72^.

### Surface plasmon resonance

Assessment of the kinetic binding parameters was performed via surface plasmon resonance (SPR) in a Biacore X-100 system. Using an adapted protocol^73^, BSA decorated with Forssman pentaose (∼76 kDa) as the ligand was prepared via reductive amination (see Fig. S3). After rebuffering into a 10 mM acetate buffer at pH 4, approximately 100 response units (RU) of ligand were immobilized onto the CM5 sensor chip surface using amine coupling as described in the manufacturer’s protocol (Cytiva). For the reference flow cell, native BSA (Sigma-Aldrich) was immobilized similarly. Purified VP1 (162 kDa) and native HPA (79 kDa, Sigma-Aldrich) buffered in SPR assay buffer (20 mM HEPES pH 7.4, 150 mM NaCl, 0.005 % Polysorbate 20) were used as analytes in different concentrations between 0.2 nM and 20 µM. All measurements were performed at 25 °C. During binding (120 s) and dissociation (180 s), a flow rate of 30 µL/min was applied. After each cycle, the surfaces were regenerated applying 10 mM glycine pH 1.5 for 120 s. Experimental curves were interpreted using BIAevaluation software (Cytiva).

### Crystallization and data collection

Purified ShPyV VP1 was concentrated to 3.5 mg/ml. Crystallization droplets comprised 1 µl protein and 1 µl reservoir solution (150 mM KSCN and 20 % (w/v) PEG 3,350). The droplets were equilibrated against the 500 µl reservoir using the hanging drop vapor diffusion method at 20 °C. For ligand derivatization, VP1 crystals were soaked in crystallization solution supplemented with the respective oligosaccharides (10 mM Forssman pentaose (F_P_), 10 mM globo-*N*-tetraose, 5 mM F_P_ + 20 mM 3’SLN, 5 mM F_P_ + 20 mM 6’SLN, 20 mM A antigen trisaccharide (all purchased from Biosynth, UK), and 20 mM Tf antigen (TCI Europe), at 20 °C for half an hour. Prior to flash freezing, crystals were transferred to oligosaccharide soaking solutions supplemented with 20 % (v/v) MPD for cryo-protection. X-ray diffraction experiments were performed at the Swiss Light Source (SLS) beamline X06DA (PSI in Villigen, Switzerland). Data were collected at a wavelength of 1 Å with an excitation time and angle of 0.1 s and 0.1° per image.

### Structure determination and refinement

X-ray diffraction images were integrated and scaled using *XDS*^74^. For each data set, 2,000-3,000 reflections were reserved as a test set for structure validation. Phases were determined by molecular replacement using the native sheep polyomavirus VP1 structure as search model (PDB 6Y61) in Phaser (CCP4)^75^. Phase refinement and model building were performed until validation parameters converged using Phenix and Refmac5, and Coot, respectively^76–78^. Oligosaccharides conformations are in accordance with the restraints from the CCP4 libraries. All protein structure representations were rendered in PyMOL (Schrödinger).

## Data availability

Atomic coordinates of ShPyV VP1 in complex with carbohydrate ligands are deposited in the RCSB Protein Data Bank (PDB) with the accession codes 7B6S (Forssman pentaose, F_P_), 7B6T (globo-*N*-tetraose), 7B6U (F_P_ + 6’SLN), and 7B6V (F_P_ + 3’SLN). Other data can be provided by the authors upon request.

## Acknowledgements

The glycan microarray studies were performed in the Carbohydrate Microarray Facility at the Imperial College Glycosciences Laboratory, which is supported by Wellcome Trust biomedical resource grants (099197/Z/12/Z, 108430/Z/15/Z, and 218304/Z/19/Z) and in part by the March of Dimes Prematurity research center grant (22-FY18-82). The sequence defined glycan microarrays contain many saccharides provided by collaborators whom we thank, as well as members of the Glycosciences Laboratory for their contribution in the establishment of the NGL-based microarray system. We also thank the German Research Foundation for funding (FOR2327 – VIROCARB, Project no. 269564371). Crystallographic data collection was performed at the macromolecular beam lines X06SA and X06DA of the Swiss Light Source, access to which was generously granted by the Paul Scherrer Institute in Villigen, Switzerland.

We thank the Max-Planck Institute for Biology in Tübingen for access to the NMR facility and Dr. Vincent Truffault for his assistance in NMR data acquisition. We also thank Prof. Barbara Mulloy (Imperial College London) for critical review of the manuscript. We thank the lab of Prof. Schulze-Osthoff at the University of Tübingen for the access to the flow cytometry appliances and especially Dr. Anja Schmitt for her help with the experiments.

## Author information

Authors and affiliations

**Interfaculty Institute for Biochemistry, University of Tübingen, Tübingen, Germany** Nils H. Rustmeier, Alexander Herrmann, Joshua C. Müller, Thilo Stehle

**Glycosciences Laboratory, Faculty of Medicine, Imperial College London, London, UK** Lisete M. Silva, Antonio Di Maio, Ten Feizi, Yan Liu

## Contributions

N.H.R. and T.S. conceptualized the study. A.H., J.C.M., and N.H.R. prepared and crystallized the protein. L.M.S and A.D.M. performed the glycan array screening experiments. A.D.M., Y.L. and T.F. performed the glycan array analyses and interpreted the glycan interaction data. N.H.R. performed and analyzed the NMR, SPR and flow cytometry experiments. A.H., J.M., and N.H.R. collected the crystallographic data and solved the structures. N.H.R. prepared the first draft of the manuscript and all contributed to the final revision. T.S. provided funding. All authors approved publication of the manuscript.

## Ethics declarations

The authors declare no competing interests. Animal blood was acquired from vendors guaranteeing animal wellbeing. Human erythrocytes were derived from anonymous donors consenting research purposes.

## Notes

### Competing Interest Statement

The authors have declared no competing interest.

### Summary of Updates

Smaller literal corrections without contentual relevance were applied to the manuscript and supplemental material S3-S6.

