## Supplemental Table S2 for "A novel and highly specific Forssman antigen-binding protein from sheep polyomavirus"

**Supplemental Table SX. Supplementary glycan microarray document based on MIRAGE Glycan Microarray guidelines (doi:**[**10.3762/mirage.3**](http://www.beilstein-institut.de/en/projects/mirage/guidelines#glycan_microarrays)**).**

| **Classification** | **Guidelines** |
| --- | --- |
| 1. **Sample: Glycan Binding Sample** | |
| Description of Sample | Sample names:  N-terminal His_6_-tagged ShPyV VP1 (ShPyV VP1-His)  Origin: recombinant  Method of preparation:  Please see the “Protein expression and purification” section under *Materials and Methods* in the main text. |
| Sample modifications | Not relevant. |
| Assay protocol | Microarray analyses were performed essentially as described ([Liu et al., Methods Mol. Biol. 2012](https://www.ncbi.nlm.nih.gov/pubmed/22057521)), for modifications of the protocol please see “Glycan microarray” under *Materials and* *Methods* section in the main text. |
| **2.** **Glycan Library** | |
| Glycan description for defined glycans | A broad-spectrum screening microarray contained 672 sequence-defined lipid-linked oligosaccharide probes, glycolipids or neoglycolipids (NGLs) was used. The probe names and corresponding structures are in **Table S1**. These are a sub-set of a recently generated large screening microarray containing around 900 glycan probes (in-house designation ‘Array Sets 42-56’, which will be published elsewhere).  The NGL probes are from the collection assembled in the course of research in the Glycosciences Laboratory (<https://glycosciences.med.ic.ac.uk/glycanLibraryList.html>).  GlyTouGan IDs (IDs in the international glycan structure repository GlyTouCan https://glytoucan.org/) are displayed for 626 out of the 672 probes in **Table S1**. |
| Glycan description for undefined glycans | Not relevant. |
| Glycan modifications | No modification was carried out for natural glycolipids.  For NGLs, unless otherwise specified these were prepared from reducing oligosaccharides by reductive amination with the amino lipid, 1,2-dihexadecyl-*sn*-glycero-3-phosphoethanolamine [(DHPE) [(Chai et al., Methods Enzymol. 2003)](https://www.ncbi.nlm.nih.gov/pubmed/12968363)]; AO, NGLs prepared from reducing oligosaccharides by oxime ligation with an aminooxy functionalized DHPE [(AOPE) [(Liu et al., Chem. Biol. 2007)](https://www.ncbi.nlm.nih.gov/pubmed/17656321)].  For full description on the definition of lipid moieties of the glycan probes please see <https://glycosciences.med.ic.ac.uk/docs/lipids.pdf>. |
| 1. **3.** **Printing Surface; e.g., Microarray Slide** | |
| Description of surface | Nitrocellulose-coated glass microarray slides. |
| Manufacturer | 16-pad UniSart® 3D Microarray Slide from Sartorius (Goettingen, Germany) |
| Custom preparation of surface | Not relevant. |
| Non-covalent Immobilisation | The lipid-linked oligosaccharide probes were formulated as liposomes by adding carrier lipids, 1,2-dihexanoyl-*sn*-glycero-3-phosphocholine (DHPC) and cholesterol for arraying and non-covalent immobilization on nitrocellulose-coated glass slides ([Liu et al., Methods Mol. Biol. 2012](https://www.ncbi.nlm.nih.gov/pubmed/22057521)). |
| **4. Arrayer (Printer)** | |
| Description of Arrayer | Nano-Plotter 2.1 (GeSiM, Radeberg, Germany). |
| Dispensing mechanism | Non-contact liquid delivery with four dispensing tips. |
| Glycan deposition | Approximately 0.33 nl was printed per spot.  Lipid-linked glycan probes were printed at 2 and 5 fmol per spot in duplicate. |
| Printing conditions | The printing solutions were all aqueous based. Printing was performed at ambient temperature and relative humidity of 58%.  The ‘liposome’ printing solutions contained 100 pmol/μl of DHPC and cholesterol (both from SIGMA) as lipid carriers in addition to the lipid-linked glycan probes. The concentrations of the lipid-linked glycan probes were 5 and 15 pmol/μl for the 2 and 5 fmol per spot levels, respectively.  The printing solutions also contained Cy3 NHS ester (GE Healthcare) at 20 ng/ml (26 fmol/μl) as a marker to monitor the printing process. |
| 2. **5.** **Glycan Microarray with “Map”** | |
| Array layout | The 672 lipid-linked probes in the screening arrays were printed on multiple subarrays for parallel binding analyses. Each pad was set up for printing 64 probes maximum, each at 2 levels in duplicate (four spots for one probe in a row); up to 256 spots (16x16) in total in each pad. |
| Glycan identification and quality control | The quality control of the screening microarrays of sequence-defined glycan probes was carried out with a panel of biotinylated plant lectins (Vector Laboratories), e.g. *Ricinus Communis* Agglutinin I (RCA_120_), *Aleuria aurantia* lectin (AAL), Concanavalin A (ConA) and wheat germ agglutinin (WGA), a wide range of anti-carbohydrate antibodies, and several commercial bacterial adhesins and toxins. Also analysed are a number of viral adhesive proteins that have been published previously, including VP1 proteins of polyomaviruses, simian virus 40 ([Campanero-Rhodes MA, et al, 2007](https://pubmed.ncbi.nlm.nih.gov/20023649/)), human JC polyomavirus ([Neu U, et al, 2010](https://pubmed.ncbi.nlm.nih.gov/20951965/)) and BK polyomavirus ([Neu U, et al, 2013](https://pubmed.ncbi.nlm.nih.gov/24130487/)), human adenovirus 52 fiber knob ([Lenman A, et al. 2018](https://pubmed.ncbi.nlm.nih.gov/29674446/)), and Chikungunya virus ([McAllister N, et al. 2020](https://pubmed.ncbi.nlm.nih.gov/32999033/)).  These data will be described elsewhere and are available upon request. |
| 3. **6. Detector and Data Processing** | |
| Scanning hardware | GenePix 4300A (Molecular Devices, UK) |
| Scanner settings | Scanning resolution: 10 μm / pixel  Laser channel: Red (scan wavelength 635 nm)  PMT: 350  Scan power: 100% |
| Image analysis software | GenePix® Pro 7 (Molecular Devices) |
| Data processing | The gpr files were entered into an in-house microarray database using software (designed by Mark Stoll, <http://www.beilstein-institut.de/en/publications/proceedings/glyco-2009>) for data processing. No particular normalization method or statistical analysis was used for the results of the screening arrays. |
| **7.** **Glycan Microarray Data Presentation** | |
| Data presentation | The microarray binding results of ShPyV-His are presented as histogram charts in **Figure 1**. The results table with binding scores and relatively binding intensities shown as ‘matrix’ are in **Table S1.** |
| 4. **8.** **Interpretation and** **Conclusion from Microarray Data** | |
| Data interpretation | No software or algorithms were used to interpret processed data. |
| Conclusions | ShPyV VP1 bound strongly with remarkable selectivity to Forssman glycolipid in the array. Little or no binding was observed to other glycan probes expect for the weak binding to α2,6-sialyl N-acetyllactosamine with a 9-O-acetylated N-acetylneuraminic acid residue. |
