## Supplemental Material S3-S6 for "A novel and highly specific Forssman antigen-binding protein from sheep polyomavirus"

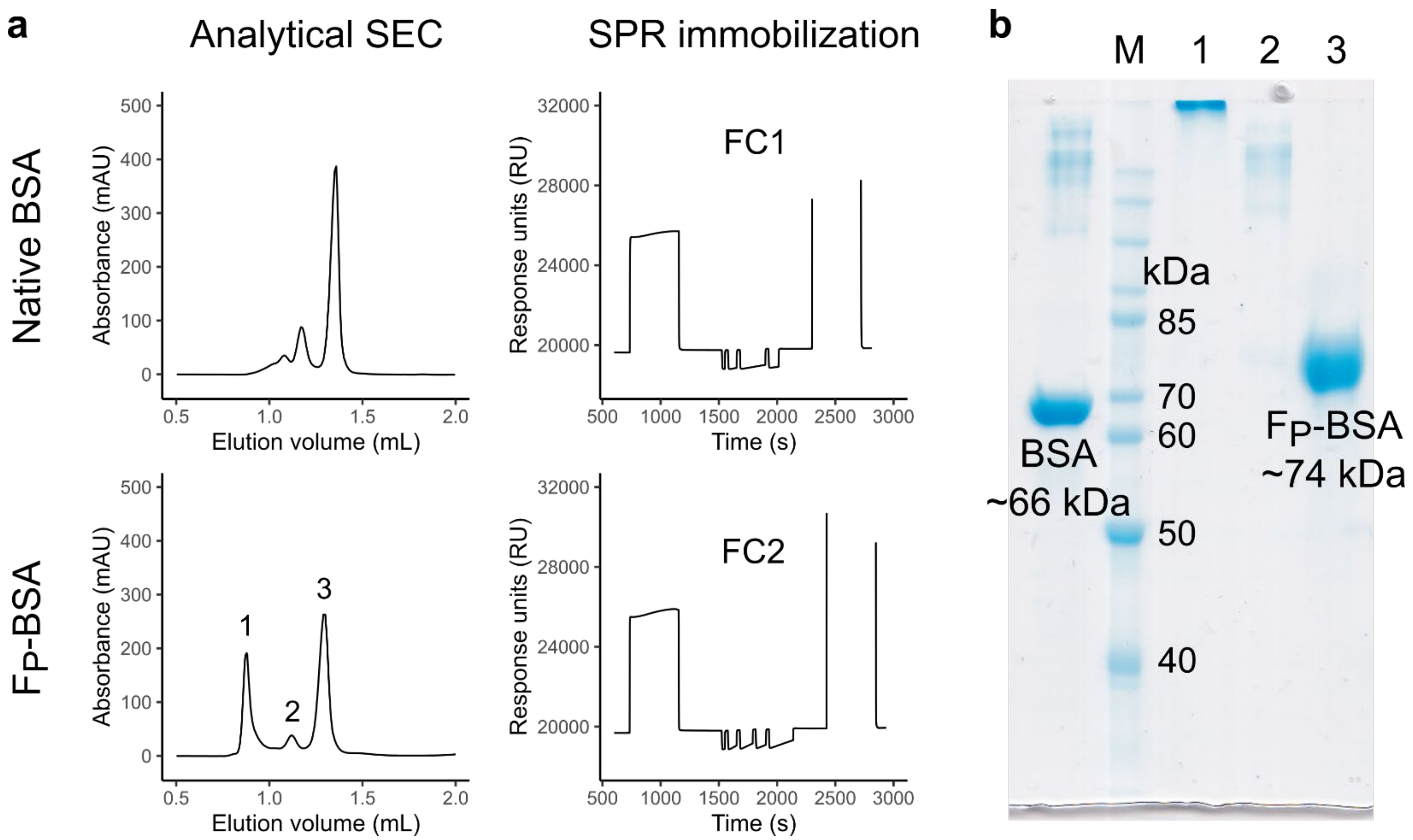


**Figure S3 a** The left panels show the analytical size exclusion chromatography (SEC) results of native BSA and Fp-BSA using a SD200 Increase 3.2/300 column (Cytiva). The right panels display the respective surface plasmon resonance (SPR) immobilization procedures. For the second flow cell (FC2), fraction 3 of the Fp-BSA SEC was used. **b** SDS-PAGE of native BSA (left lane) and the three fractions of the analytical SEC of Fp-BSA. The mass difference between BSA and Fp-BSA (lane 3) corresponds to a coupling of ~8 molecules of Forssman pentaose to one molecule of BSA.

**Table S4** Data collection and refinement statistics for the glycan complex crystal structures of ShPyV VP1.

| Data set of derivate(s) | **Forssman pentaose** | **Globo-*N*-tetraose** | **Forssman pentaose + 6'SLN** | **Forssman pentaose + 3'SLN** |
|---|---|---|---|---|
| PDB accession code | 7B6S | 7B6T | 7B6U | 7B6V |
| **Data collection** | | | | |
| Space group | $P3_1$ | $P3_1$ | $P3_1$ | $P3_1$ |
| a, b, c (Å) | 130.5, 130.5, 222.4 | 130.3, 130.3, 222.2 | 129.6, 129.6, 220.4 | 130.4, 130.4, 221.2 |
| α, β, γ (°) | 90, 90, 120 | 90, 90, 120 | 90, 90, 120 | 90, 90, 120 |
| Resolution (Å) | 42.71 - 1.92 (2.04 - 1.92) | 48.91 - 1.70 (1.80 - 1.70) | 49.47 - 1.90 (2.02 - 1.90) | 48.85 - 1.80 (1.91 - 1.80) |
| Total reflections | 2'214'763 | 2'676'281 | 1'728'348 | 3'864'575 |
| Unique reflections | 321'069 | 462'792 | 325'151 | 390'469 |
| $R_{meas}$ (%) | 9.3 (68.4) | 10.7 (108.5) | 14.6 (122.3) | 16.2 (127.3) |
| I/σI | 16.67 (2.91) | 13.51 (1.61) | 9.46 (1.33) | 11.02 (1.51) |
| $CC_{1/2}$ (%) | 99.9 (78.6) | 99.8 (48.5) | 99.6 (44.1) | 99.7 (47.1) |
| Completeness (%) | 99.8 (98.7) | 99.9 (99.7) | 99.8 (98.7) | 99.8 (98.9) |
| Wilson B-factor ($Å^2$) | 32.9 | 27.9 | 31.3 | 28.5 |
| **Refinement** | | | | |
| $R_{work}$ / $R_{free}$ (%) | 14.9 / 17.6 | 14.3 / 16.5 | 14.6 / 17.3 | 15.2 / 17.7 |
| No. of atoms | | | | |
| Protein | 20'468 | 20'343 | 20'393 | 20'322 |
| Water | 2'254 | 3'256 | 2'031 | 2'373 |
| Glycan | 564 | 411 | 761 | 636 |
| B-factor ($Å^2$) | | | | |
| Protein | 29.5 | 22.8 | 28.0 | 24.5 |
| Water | 37.4 | 34.8 | 35.8 | 30.8 |
| Glycan | 32.8 | 37.2 | 34.0 | 26.4 |
| R.m.s.d | | | | |
| Bond length (Å) | 0.0093 | 0.0095 | 0.0090 | 0.0092 |
| Bond angle (°) | 1.65 | 1.64 | 1.65 | 1.72 |
| Ramachandran outliers (%) | 0.08 | 0.08 | 0.04 | 0.15 |
| Clashscore | 1.94 | 1.80 | 1.89 | 2.96 |

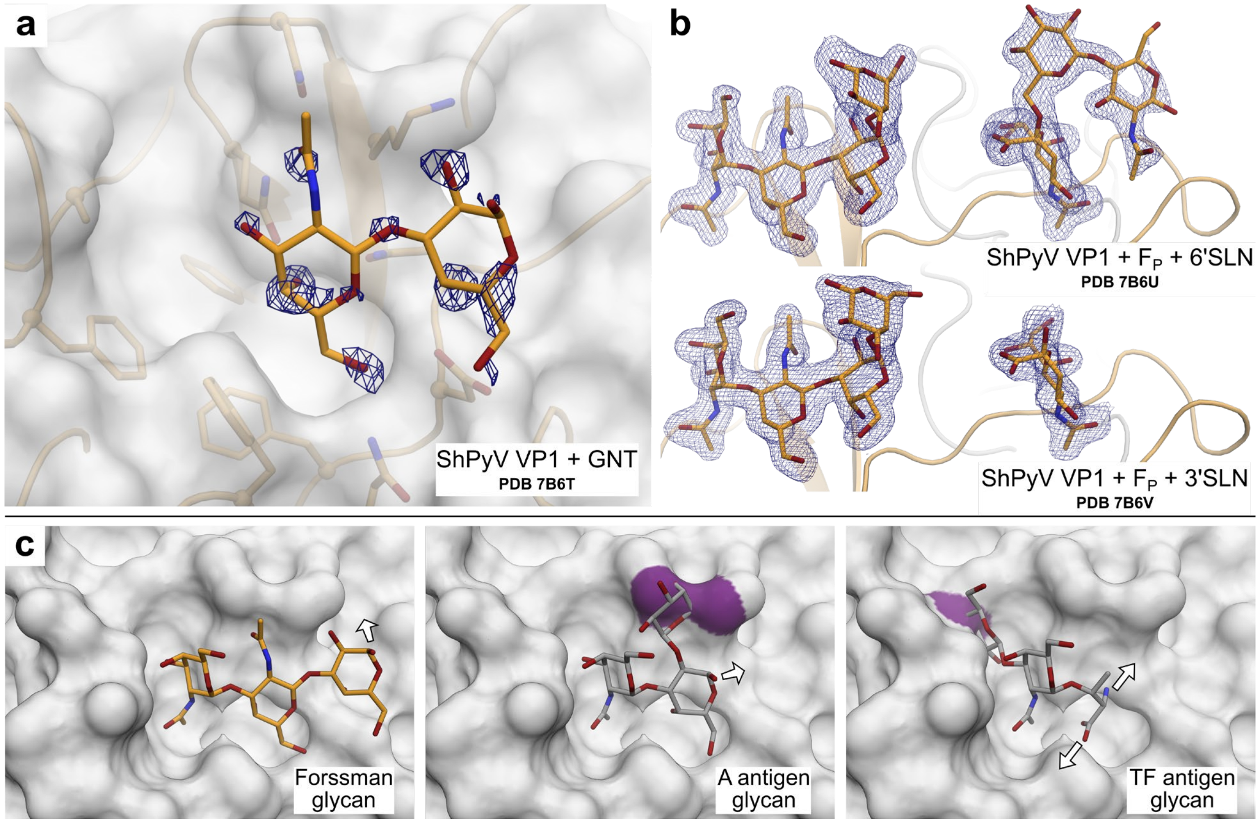


**Figure S5 a** The crystal structure of ShPyV VP1 in complex with 10 mM globo-*N*-tetraose (GNT, PDB 7B6T), the glycan of the Forssman precursor molecule globoside, reveals only negligible binding. **b** The co-complexes of ShPyV VP1 with Forssman pentaose ($F_P$), and 6'sialyllactosamine (6'SLN, PDB ID 7B6U) or 3'sialyllactosamine (3'SLN, PDB ID 7B6V) show that the different glycan species bind to different sites on the protein surface with no close contact between them. In **a** + **b**, the Fo-Fc electron densities are shown as blue meshes around the individual oligosaccharides with a contour level of 2.5 σ and a radius of 1.6 Å. **c** Based on αGalNAc superposition, the side-by-side depiction of the Forssman antigen glycan (GalNAcα1-3GalNacβ1-3Galα), A antigen glycan (GalNAcα1-3[Fucα1-2]Galβ), and the TF antigen (Galβ1-3GalNAcα1-Ser/Thr) in association with the VP1 surface. In the A antigen and TF antigen glycan models the fucose and galactose, respectively, result in overlaps with the protein (highlighted in purple). The Tn antigen, which corresponds to TF truncated by the terminal galactose, also cannot bind to VP1 as the glycoprotein proceeds horizontally to VP1, which again would result in overlaps. In all panels of **c**, arrows indicate the linkage to the remaining antigen portion. Throughout the figure, oxygen and nitrogen atoms are colored red and blue, respectively.

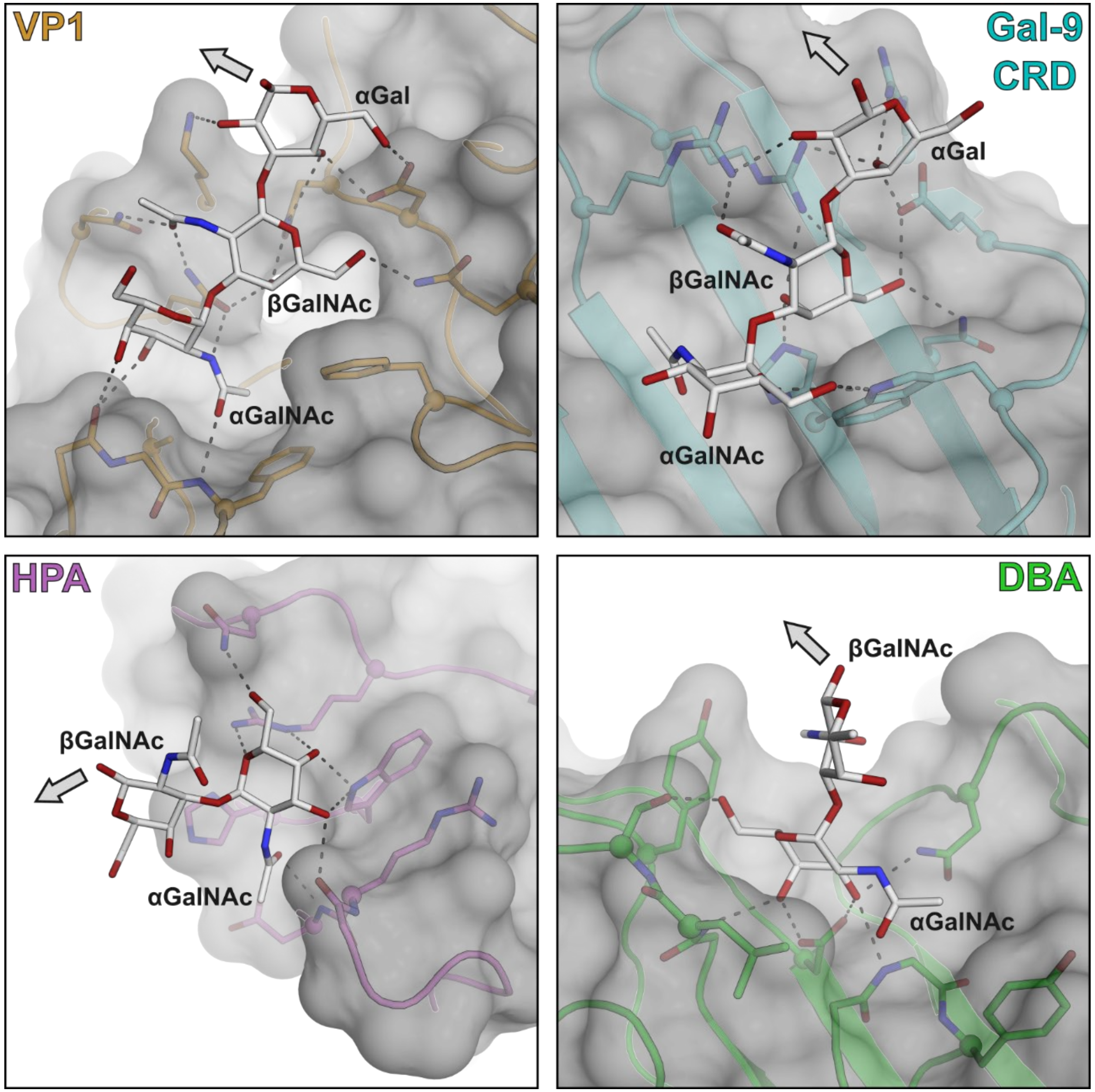


**Figure S6** Different modes of Forssman glycan binding in ShPyV VP1, Galectin-9 carbohydrate recognition domain (Gal-9 CRD), *Helix pomatia* agglutinin (HPA), and *Dolichos biflorus* agglutinin (DBA). VP1 recognizes the GalNAcα1-3GalNAcβ1-3Galα trisaccharide sequence of Forssman pentaose ($F_P$) via multiple hydrogen bonds to each sugar unit (upper left panel). Human Gal-9 CRD (PDB 5X4A) mainly interacts with the βGalNAc of $F_P$ (upper right panel). HPA (PDB 2CGY) and DBA (PDB 1LU1) interact exclusively with the terminal αGalNAc but not βGalNAc (bottom panels). All proteins are shown as transparent surfaces with an underlying cartoon representation. Binding site residues (colored according to the protein) and Forssman glycans (light grey) are shown as sticks. Arrows indicate the direction of the linkage to the next monosaccharide. Oxygen and nitrogen atoms are colored red and blue, respectively. Hydrogen bonds are indicated as dashed black lines.

Forssman glycan binding by *Sinularia lochmodes* lectin-2 (PDB 5X4A) is homologous to HPA and not shown.
